# Islands of hope in the Sahel: farmer-managed grazing exclusions restore biodiversity and boost livelihoods

**DOI:** 10.64898/2026.01.16.699859

**Authors:** Gabriel Marcacci, Roland R. Kaboré, Ambroise N. Zongo, Serge T. Zoubga, Bakary Diakité, Reto Spaar, Franziska Kaguembèga-Müller, Alain Jacot

## Abstract

Global restoration initiatives to address climate change, biodiversity loss, and land degradation often remain fragmented, limiting opportunities for synergistic outcomes. Cost-effective bottom-up approaches, such as participatory community-led natural regeneration, have substantial potential but are often overlooked, hindering their wider implementation. In drylands such as the Sahel, where more than half of rangelands are degraded, restoration is urgently needed yet remains understudied. We evaluated a participatory model based on farmer-managed grazing exclusions (≈3 ha each) established by local communities in Burkina Faso. Using paired comparisons of 54 exclusions and adjacent controls, we combined vegetation and bird surveys with socioeconomic interviews to quantify ecological recovery and livelihood impacts.

Grazing exclusions substantially enhanced biodiversity, with tree richness and abundance 122% and 362% higher, respectively, and bird richness 20% higher than controls. Carbon sequestration increased by 18%, vegetation productivity by 43.9% and ecosystem services multifunctionality by 282%. Farming households managing exclusions reported more than double the annual income of those without (+115.2%), with 28.5% of economic gains directly attributable to harvested natural products derived from ecosystem services provided by restored vegetation. Grazing exclusions increased household income both directly and indirectly through their effects on tree richness and ecosystem services multifunctionality. These findings demonstrate that community-led natural regeneration through small-scale grazing exclusions is a cost-effective, multifunctional nature-based solution that simultaneously restores biodiversity, enhances ecosystem services, and supports livelihoods. Scaling this approach could substantially advance global biodiversity, climate, and sustainable development goals, but will require governance and financing mechanisms that recognize the value of bottom-up initiatives.

**Significance Statement:** Dryland ecosystems such as the Sahel are among the most extensive and degraded landscapes on Earth, yet they remain underrepresented in global restoration efforts. We show that farmer-managed grazing exclusions – small areas where livestock are excluded to allow natural regeneration of the vegetation – can simultaneously restore biodiversity, enhance ecosystem services, and double household income. This bottom-up, community-led approach delivers synergistic outcomes, highlighting the power of natural regeneration as a cost-effective restoration strategy. By linking ecological recovery with tangible livelihood benefits, grazing exclusions offer a scalable model that advances global biodiversity, climate, and development goals. Supporting such locally grounded initiatives will be essential for building resilient and equitable landscapes worldwide.

## Introduction

Anthropogenic activities are driving our planet beyond safe biophysical limits, with six out of nine planetary boundaries already transgressed (1). The multiplication of global crises such as climate change, biodiversity loss and ecosystem degradation threaten the functioning of ecological systems and the livelihood of communities that depend upon them (2). In response, global assessments (e.g., IPBES, ICCP) and international conventions have set ambitious targets such as the Paris Agreement for climate mitigation, Kunming-Montreal Global Biodiversity Framework for biodiversity conservation, the Sustainable Development Goals to improve human well-being, and the Land Degradation Neutrality targets or the UN Decade on Ecosystem Restoration to halt and reverse land degradation. Yet, despite recognition that these crises are interlinked, solutions often remain fragmented – undermining the potential for synergistic outcomes (3–4). For example, large-scale habitat restoration initiatives often prioritize mass tree planting to sequester carbon or halt desertification at the expense of comparatively greater ecological and socio-economic benefits offered by cost-effective natural forest regeneration (5–8). Such large initiatives are attractive as they define clear targets (e.g., planting one billion trees) which are politically appealing and easily communicable (9). But they risk overlooking more holistic, bottom-up land-use approaches that simultaneously mitigate climate change, restore biodiversity, enhance ecosystem services, and support local livelihoods (3, 10–12).

While most of the pledges have been directed to restoration of tropical forests, ecosystems such as arid woody savannah or rangelands have received far less attention (3). Yet, arid rangelands such as the Sahel Zone (a vast area spanning 12 countries and home to over 400 million people; International Fund for Agricultural Development www.ifad.org) in sub-Saharan Africa are the most extensive use of land worldwide, and more than half of them are already degraded (12–14). Land degradation here is driven primarily by overgrazing by free-ranging domestic livestock and unsustainable use of natural resources, especially deforestation for fuelwood and charcoal production (13, 15–17). These pressures not only erode biodiversity but also undermine ecosystem services critical to rural livelihoods. In turn, land degradation is responsible for increasing food insecurity and poverty, threatening already vulnerable human populations (17–18).

To halt and reverse land degradation in the Sahel Zone and mitigate climate change, large-scale initiatives – such as the Great Green Wall (a continental “green belt” from Senegal to Djibouti) and AFR100 (a pledge to restore 100 Mha in Africa by 2030 under the Bonn Challenge) – represent ambitious responses to this crisis and have mobilized substantial political and financial capital (17, 19). But more than half of the pledged area is set to become plantations of commercial trees and only one-third under natural regeneration despite it is often the most cost-effective option (5–6) – although certain conditions need to be met to favor natural regeneration (10). To increase the ecological resilience and the provision of multiple ecosystem services that support livelihoods we need multifunctional and species diverse restored forests (10, 20). Moreover, while essential for raising awareness and coordinating regional action, large top-down initiatives face persistent challenges, including land tenure insecurity, sociopolitical instability, and limited integration of biodiversity and local community needs (3,12, 21). Achieving global biodiversity, climate, and sustainable development targets will require complementing top-down strategies with locally grounded, bottom-up “grass-root” approaches that are adaptive, inclusive, and cost-effective (11). Yet, despite their promise, they remain understudied. More scientific evidence is needed to understand how bottom-up and participatory approaches to natural regeneration can deliver synergistic ecological and economic benefits for ecosystem restoration, biodiversity recovery and human well-being (10–12).

In this study, we focused on the work of the international organization newTree (www.newtree.org) and its local partner in Burkina Faso, the association tiipaalga (www.tiipaalga.org). Together, they have developed a holistic and participatory approach to land restoration that empowers farming households to become active agents of transformative change. At the core of their work is the establishment of small-scale grazing exclusions: areas of approximately three hectares where domestic livestock are excluded to allow natural vegetation to regenerate. This form of assisted natural regeneration is aimed to support both ecological and livelihoods recovery (10, 22–23).

Beyond land restoration, tiipaalga provides training and long-term support in sustainable land management to reduce deforestation and diversify household income through the development of income-generating activities linked to the provisioning services valued by rural communities and provided by restored ecosystems in grazing exclusions. These include for instance sustainable fuelwood harvesting through tree pruning, processing of non-timber forest products (e.g., transforming shea nuts into shea butter), hay harvesting for livestock or market sale, or the promotion of apiculture as a high-value livelihood strategy. At the same time, tree regeneration and reduced deforestation help mitigate CO _2_ emissions. This holistic approach may have a great potential to build ecological resilience and strengthen the adaptive capacity of rural communities.

Previous studies have shown that grazing exclusions can enhance tree regeneration and biodiversity recovery across various regions (24–28), including Burkina Faso (29–30), suggesting their potential as a promising land restoration strategy. Moreover, they also restore biophysical processes such as soil fertility and water retention (31) and can improve the livelihood of local communities (11, 32). However, few studies have documented their combined and interlinked ecological and economic benefits. In this unique semi-experimental study, we used a paired design (grazing exclusions are compared to paired control sites) to evaluate the multiple ecological and economic benefits provided by grazing exclusions. To do so, we conducted biodiversity surveys (birds – via passive acoustice monitoring – and vegetation – via field inventories and remote sensing) as well as socioeconomic interviews in 54 sites, from which we derived six indicators (see Table S1) of provisioning ecosystem services, which were also combined in a single metric – ecosystem services multifunctionality – following state-of-the art methodologies (33). Next, we combined biodiversity and socioeconomic data in structural equation models to assess the direct and indirect contributions of biodiversity and ecosystem services provided by grazing exclusions to rural economies, thus allowing a mechanistic understanding of the underlying effects. We expect grazing exclusions to promote biodiversity and ecosystem services multifunctionality, which in turn increase household income. We show that small-scale grazing exclusion combined with training and long-term support is a promising restoration strategy that yields synergistic outcomes including substantial ecological and economic gains, offering a scalable model to boost dryland restoration efforts that align with global sustainability goals. Yet, there exist several barriers that need to be unlocked to upscale this holistic and participatory approach of natural regeneration to other regions, for which we provide practical recommendations and suggest future research directions.

## Results

### Effect of grazing exclusion on biodiversity, ecosystem services and income

Grazing exclusion had a consistently positive impact on biodiversity, ecosystem services and annual income, underscoring its potential to deliver multiple ecological and economic benefits simultaneously (Figure 1; Table S3). Tree species richness and tree abundance were 122.5% and 361.5% higher, respectively, in grazing exclusions compared to control sites. Similarly, bird species richness increased by 20% in exclusion sites. Although CO_2_ sequestration was 18.3% higher in exclusion sites, this effect was not statistically significant. However, NDVI – a reliable proxy for primary productivity and, indirectly, CO_2_ sequestration – was statistically 43.9% higher in exclusion sites (Figure S2). Ecosystem services multifunctionality (ES-multifunctionality), which quantified the simultaneous delivery of multiple ecosystem services (apiculture, construction wood, firewood, fodder, medicinal plants, and non-timber forest products which correspond to fruits and nuts), was 282.2% higher in grazing exclusions. This positive effect was consistent across both average and threshold-based (i.e., number of ecosystem services indicators that cross a certain threshold, expressed as the percentage [1-99%] of the maximum observed values of each indicator within our study sites) multifunctionality metrics, regardless of the threshold, though the maximum differences were at intermediate thresholds (10-50%; Figure S3). Moreover, when looking at each ecosystem services individually, they are also all increased in grazing exclusions compared to control sites: +385.7 % for apiculture, +329 % for construction wood, +367.6 % for firewood, +245.9 % for fodder, + 1500 % for medicinal plants and + 180.9 % for non-timber forest products (Figure 3B, Figure S4). Finally, farming households managing grazing exclusions reported annual incomes more than twice as high as those without grazing exclusions.

**Figure 1.**
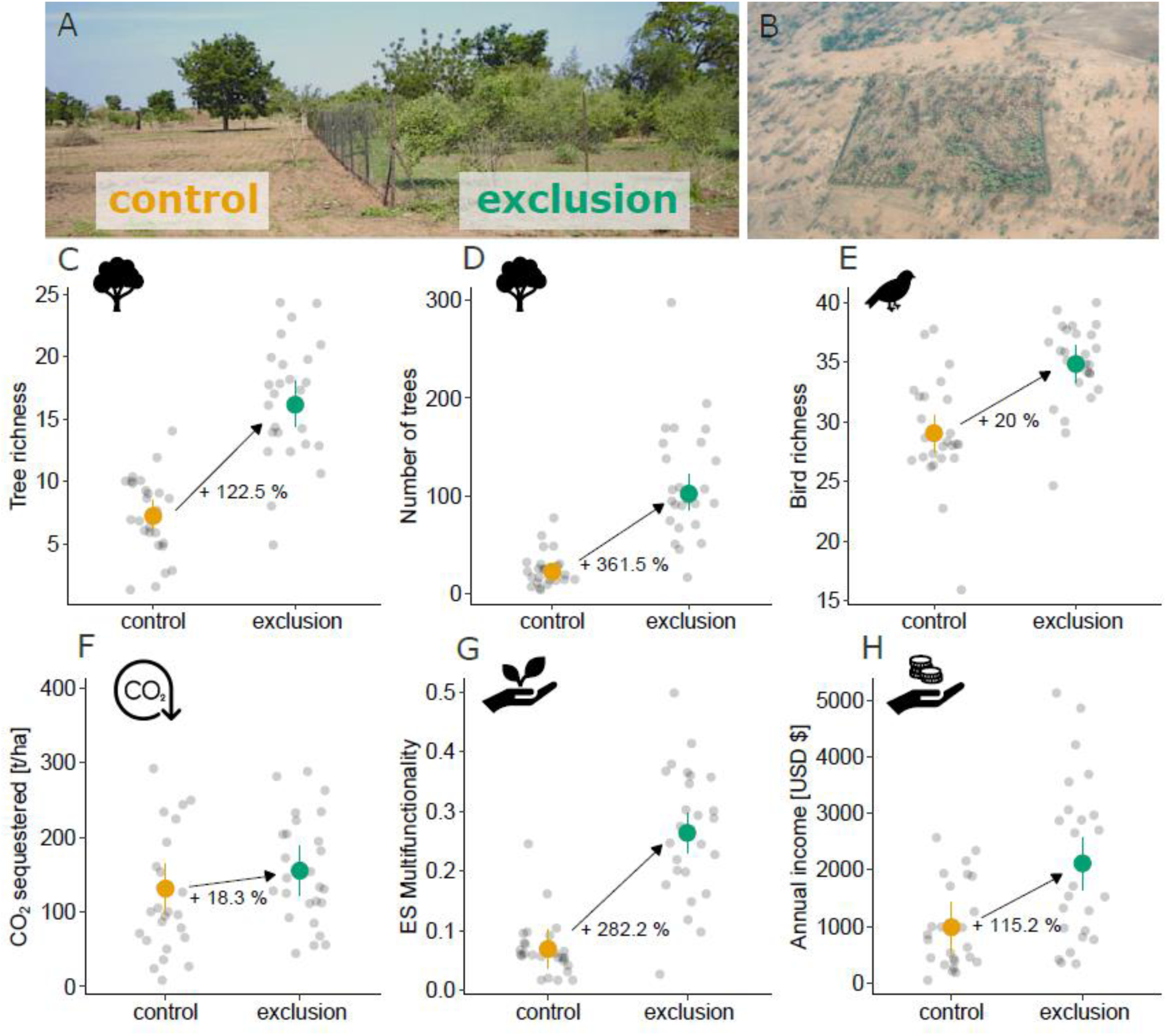
Picture of regenerated trees within the grazing exclusion (A) to the right, contrasting with degraded land outside of the grazing exclusion (control) to the left. Aerial image of a grazing exclusion (B) and its surrounding degraded landscape. Positive effect of grazing exclusions on tree richness (C), tree abundance (D), bird richness (E), CO_2_ sequestration (F), ecosystem services multifunctionality (G) and household income (H). Grey dots depict raw data, colored points (yellow for controls and green for grazing exclusions) the predicted means and confidence intervals drawn from (generalized) linear mixed-effects models. Note that annual income was converted into US dollars from CFA francs (XOF) with converting factor 1 XOF = 0.0018 USD (30.06.2025).

### Direct and indirect effects of grazing exclusion, tree richness and multifunctionality on income

We used structural equation modeling (SEM) to evaluate how grazing exclusion, time (since the establishment of grazing exclusion), and tree richness influenced ES-multifunctionality and annual income (Figure 2A). The final SEM showed good fit (Fisher’s C = 5.72, p-value = 0.455), with no evidence of significant missing paths. Tree richness was positively associated with ecosystem multifunctionality (β (unstandardized) = 0.211 ± 0.056 SE, p-value = 0.024) and indirectly increased annual income via this effect. Grazing exclusion had strong and consistent effects on all variables. Compared to controls, grazing exclusions significantly increased tree richness (β = 0.786 ± 0.090, p-value < 0.001), which in turn promoted multifunctionality. However, grazing exclusions also had a direct positive effect on annual income (β = 0.762 ± 0.226, p-value < 0.001), independent of changes in biodiversity or ecosystem services. Time had a smaller but significant effect on tree richness (β = 0.042 ± 0.190, p-value = 0.029), indicating tree richness increased with time since the establishment of a grazing exclusion. Multifunctionality, while positively associated with income (β = 0.566 ± 0.866), did not have a statistically significant direct effect on income (p-value = 0.52), though its contribution to total effects was evident through indirect paths.

**Figure 2.**
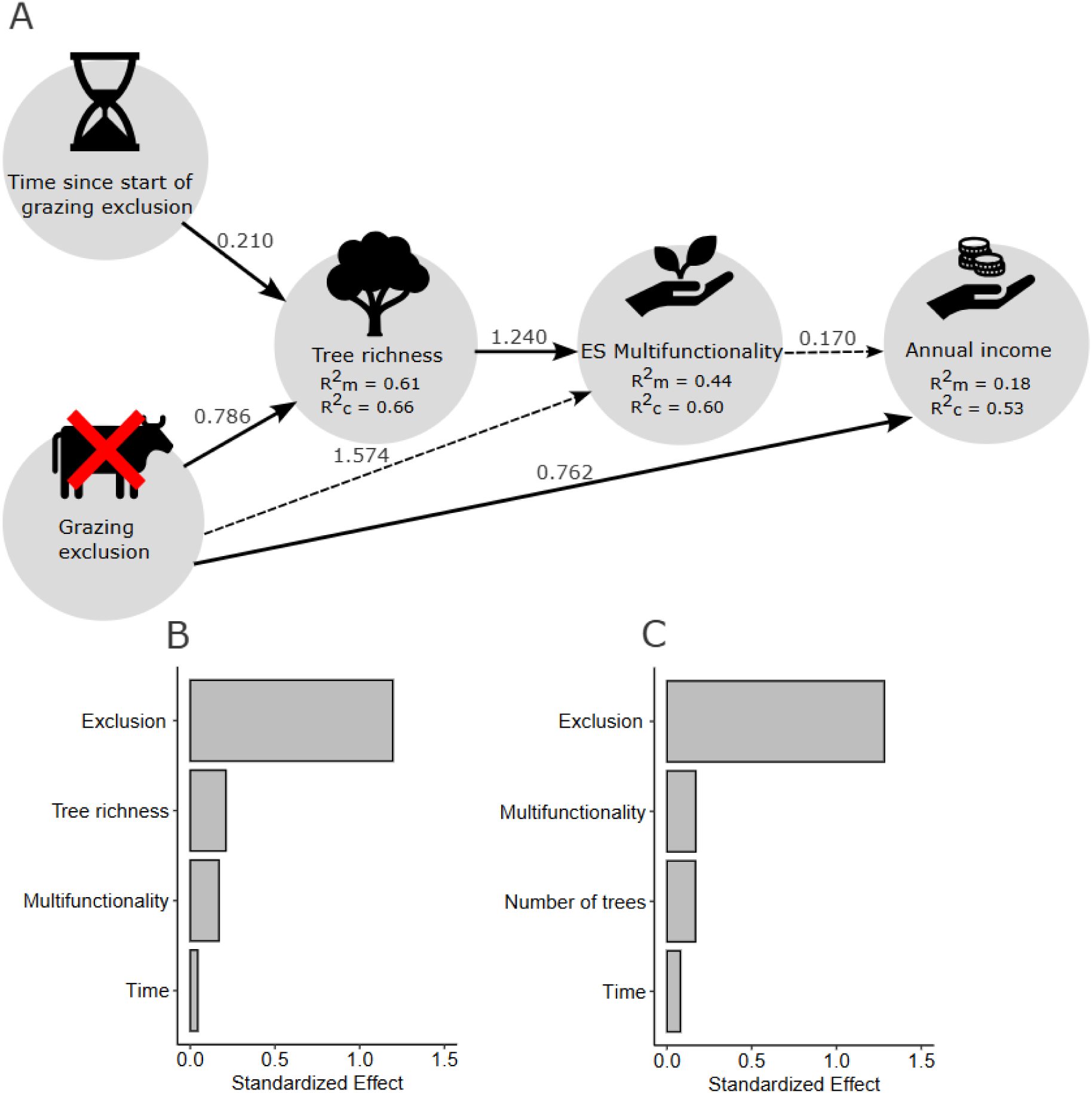
Path diagram (A) illustrating the results of a Structural Equation Model investigating the direct and indirect effects of grazing exclusions, time, tree richness and ES-multifunctionality on household income (the same path diagram but with tree abundance instead of tree richness is available in Figure S5). Significant (p-value < 0.05) causal relationships are displayed with solid arrows and non-significant relationships with dotted arrows. Standardized models’ coefficients are displayed next to the arrows and marginal and conditional R^2^ of individual model below the response variables. Bar plots showing the total (direct + indirect) standardized effect of grazing exclusion, tree richness / abundance, ES-multifunctionality and time on household income. The same SEM was fitted once with tree richness (B) and once with tree abundance (C).

To compare the relative importance of predictors, we calculated standardized total effects (direct + indirect, see Figure 2B). Grazing exclusions had the largest total effect on annual income, followed by tree richness and multifunctionality. Tree richness had a substantial indirect effect on income through its impact on multifunctionality, whereas grazing exclusion’s effect was both direct and mediated. Standardized coefficients are reported in Figure 2, and all model paths and coefficients are detailed in Table S4. The same analysis was repeated with tree abundance and tree richness, which yielded similar results (see Figure 2C, Table S5 and Figure S5).

### Sources of income

Face-to-face socioeconomic interviews revealed that farming households managing grazing exclusions reported annual incomes more than twice as high as those without access to such areas (982 *vs* 2114 USD $). On average, 28.5% of this income was directly attributable to the sale of natural resources harvested within the grazing exclusions (Figure 3A). These resources corresponded to the provisioning services assessed in our study (Figure 3B). In addition to these direct economic gains, farming households also benefited from indirect contributions of grazing exclusions to crop or livestock production that are more difficult to quantify (e.g., beyond selling fodder, many households used it to feed their own livestock, thereby enhancing livestock productivity) or the direct consumption of natural products, as they are not consistently recorded.

**Figure 3.**
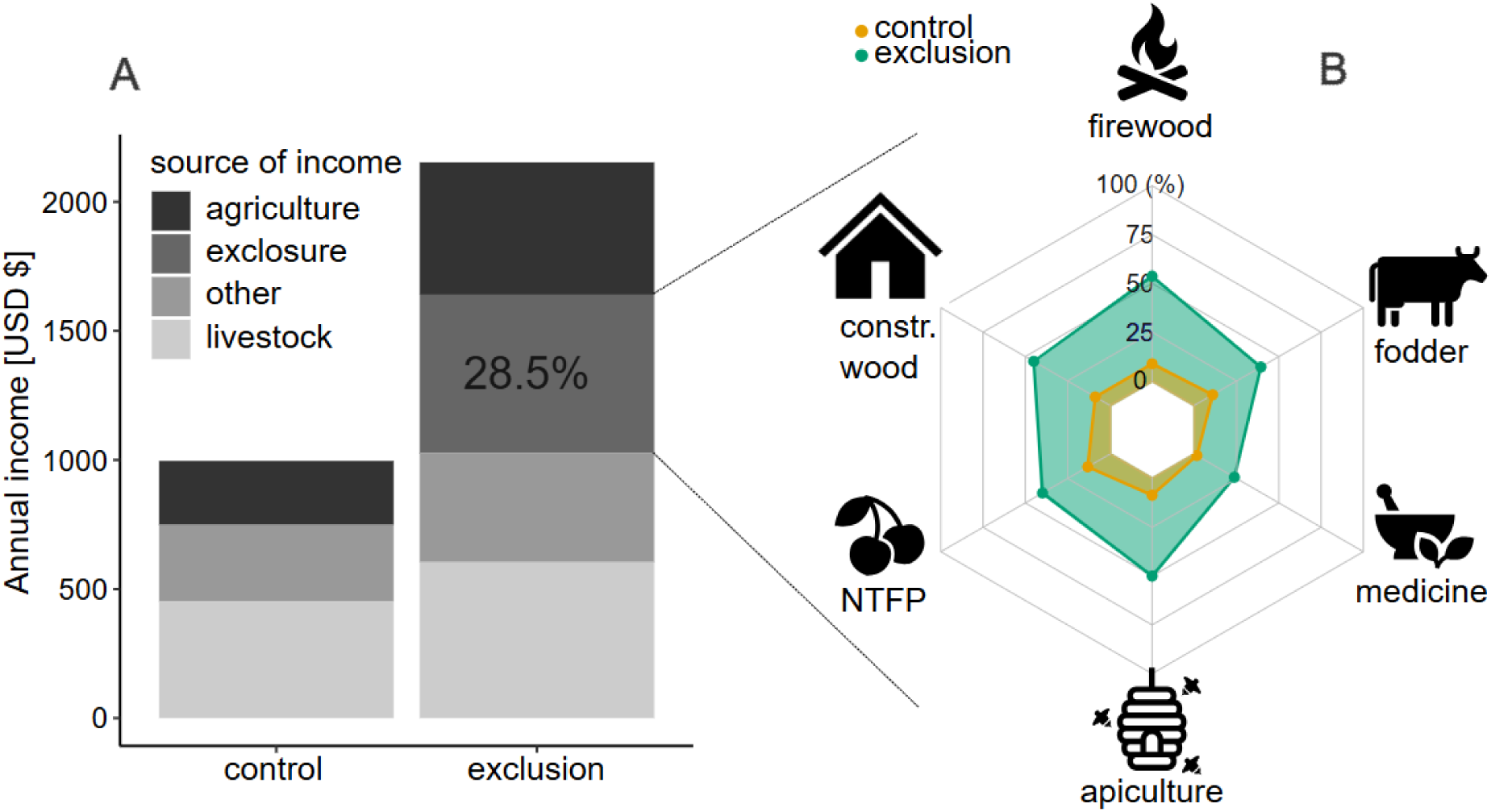
Different sources that contribute to annual household income (A) in controls and grazing exclusions: crop production (“agriculture), livestock production (“livestock”), non-agricultural activities (“other)”, and income derived from the sale of resource-based products from grazing exclusions (“exclusion”). The sale of products harvested in grazing exclusions contributes to 28.5 % of annual household income. Radar chart (B) illustrating the provisioning ecosystems services from which these products are derived, here standardized between their respective five maximum and five minimum values. Note that annual income was converted into US dollars from CFA francs (XOF) with converting factor 1 XOF = 0.0018 USD (30.06.2025).

## Discussion

### Grazing Exclusions: A Cost-Effective Solution to Multiple Crises

Our findings demonstrate that assisted natural regeneration via grazing exclusions are a model of nature-based solutions that can deliver significant ecological and economic benefits, offering a cost-effective and scalable solution to multiple, interlinked global challenges such as climate change, biodiversity loss, land degradation, and poverty. While it has already been shown that grazing exclusion has great potential to restore degraded lands (29, 34), here we showcase that such local and bottom-up initiative fosters biodiversity recovery and sustainable livelihoods (household income and food security), thus generating synergistic outcomes that align with global restoration and sustainability targets. Importantly, our study revealed the importance of indigineous tree diversity for ecosystem services provision and household income, indicating that it is critical to promote tree diversity in restoration efforts and to diversify planted forests when natural regeneration is not possible (20, 33, 35). This holistic approach supports progress toward at least 7 Sustainable Development Goals (SDGs), including SDG 1 (No Poverty), SDG 2 (Zero Hunger), SDG 13 (Climate Action), and SDG 15 (Life on Land), as well as multiple targets under the Kunming-Montreal Global Biodiversity Framework (GBF), notably Targets 2 (ecosystem restoration), 9 (benefits for people), 11 (Nature’s contributions to people), 14 (integration of biodiversity across sectors) and 22 (local communities participation).This emphasizes the critical importance of mainstreaming biodiversity in human or climate-centered restoration initiatives to meet multiple goals (3, 8).

### Upscaling this Model System to Boost Dryland Restoration Efforts

Our study highlights small-scale grazing exclusions as a replicable model for dryland restoration, but effective upscaling requires thoughtful considerations. To ensure lasting success, ecological recovery needs to be directly linked to livelihood benefits. This is where participatory approaches rooted in local governance (whether at household, cooperative, or community level) have proven more sustainable than top down interventions as it strengthens local ownership, aligns restoration outcomes with traditional knowledge systems, and guarantees that ecological and livelihood benefits are equitably shared (11). Yet, while the basic mechanism of grazing exclusions – protecting land to enable natural regeneration – is broadly transferable, its success is context-specific and depends on aligning with local socioeconomic realities and cultural practices as well as environmental and socio-ecological conditions (10, 36–37). In our study, large direct economic benefits – doubling household income – were key to the popularity of this approach and tiipaalga has the resources to consider only one quarter of the applications (to support the establishment of new grazing exclusions) they receive from local communities (29). Tree diversity had an indirect positive effect on income via its impact on multifunctionality, underscoring the critical role of ecosystem services and income-generating activities in making restoration viable and attractive (34, 38).

Although the direct effects of ecosystem services provision on household income were not significant in our analysis, we showed that grazing exclusions contributed up to almost a third of household economy thanks to the sale of natural resources harvested in these restored ecosystems. This figure is similar to others studies conducted in Burkina Faso and Northern Ghana (11, 32).This is particularly true for natural products which have a high market value such as shea butter or honey that can be an important source of income, especially for women (12, 39). Yet, aligning with Weston et al., (2015), we found that non-market benefits generated a large share of direct and indirect (not measured) economic gains (e.g., own consumption of natural products, improved crop and livestock productivity, enhanced leadership capacity to engange in other off-farm economic activities), emphasizing the need for a holistic approach. Upscaling should therefore prioritize livelihood benefits alongside restoration goals, including social consideration of local communities and empowering them (37). Without this, restoration efforts risk being short-lived or socially contested.

Nevertheless, more in-depth ecological and social studies are needed across different socio-ecological contexts to minimize the risk of negative outcomes. Indeed, modifying the complex dynamics of such social-ecological systems (e.g., arid rangelands) can have unexpected consequences, as for example livestock grazing can also have positive impact on native plant communities in some ecosystems and circumstances (40), or create conflicts among pastoralists or leading to environmental injustice depending on the types of land tenures and property rights of communal rangelands (41). Furthermore, fencing is not an appropriate measure in regions where large wild herbivores are still widely present (42).

### Barriers and Opportunities to Finance Natural Regeneration Initiatives

The cost of financing the restoration of degraded lands is estimated to amount to 18-70 billion in the Sahel alone (17), which represent a massive challenge with lack of funding one of the main barriers to restoration efforts (43). This is especially true in recent times, with public funding for environmental and development aids massively reduced or cut by politic administrations worldwide. While most restoration projects are still funded by public institutions, there is a growing interest in private funding (44). Yet, whether it is through CO_2_ credits or other types of investments, private investors often see restoration projects as high-risk investments with a low return (9). Moreover, private investors tend to exclude investments in “high-risk areas” (e.g., politically unstable countries from the Sahel zone, which are often the ones in greater need of support for restoration projects) and projects focusing on natural regeneration because of the lack of clear and easily communicable targets and achievements in contrast to tree planting (9). Moreover, small grass-root and community-led efforts often struggle to get funding due to low visibility. These represent major barriers to finance local natural regeneration initiatives like farmer-managed small-scale grazing exclusions, despite their cost-effectiveness (11). As one of the main benefits for private investors to finance restoration projects is improving their brand image, it is crucial to better communicate the large and multiple benefits (beyond the area of land restored) provided by these small-scale natural regeneration initiatives, including the results of scientific publications, to different stakeholders in the private sector and the wider public. In addition, more research should explore the potential opportunities for using the carbon market to fund a wider variety of restoration initiatives that provide ecological and social benefits. Finally, this also emphasizes the need for robust and transparent monitoring and reporting systems that are scalable to projects of different sizes and forms to ensure that the targets and promises translate into measurable outcomes that really make an impact (11, 45).

### Study limitations and Future Research Directions

While our study demonstrates clear ecological and economic benefits of small-scale grazing exclusions, it has several limitations. First, our study area is located in the Sahel region, which sadly suffers from insecurity, limiting access to the study sites, with concomitant impact on fieldwork activities and research possibilities. Nevertheless, the Sahel region extends from Senegal to Djibouti, and remains poorly studied, making our study highly valuable as restoration efforts in the region are urgent. Second, the benefit of grazing exclusions may not be immediate and only manifest after a few years, which can affect local acceptance. Although all sites in our study were at least six years old, and we found little effects of time, longer-term dynamics remain unknown. For example, benefits may plateau or decline if certain plant species become dominant (46, 47).

Indeed, as savannahs are dynamic ecosystems shaped by herbivores, some levels of control grazing could enhance long-term sustainability (46, 47). Third, the grazing exclusions studied were implemented by NGOs under real-world conditions rather than experimental designs. While this increases relevance for applied restoration, it limits our ability to test variables such as size of the exclusions, their spatial arrangement, the effects of tree identity or adaptive grazing regimes. A controlled restoration experiments, such as tree biodiversity experiments (e.g., Tree Diversity Network (48)) would be valuable to answer these questions. Fourth, we did not measure regenerating trees, which yields underestimation of CO_2_ sequestered in grazing exclusions. Following established and standardized protocols that included regenerating trees (e.g., SEOSAW protocol (49)) would improve these estimations and possibly increase the interest of investors. Lastly, our study focused primarily on provisioning services and household income. However, restored ecosystems also provide regulating and supporting services (e.g., pollination, pest control, water retention, and cultural values), which warrant further investigation. Notably, local communities expressed keen interest in the return of wildlife such as birds, underscoring the importance of incorporating local perceptions into restoration planning to foster long-term support and engagement and to prevent human-wildlife conflicts.

## Conclusion

Our study revealed the great potential of assisted natural regeneration via small-scale grazing exclusions as a cost-effective nature-based solution which can restore degraded landscapes in multifunctional ecosystems which enhance biodiversity and living conditions of local communities. This holistic and participatory approach to restoration delivers synergistic ecological and economic outcomes that align with global sustainability targets, including the SDGs and the Global Biodiversity Framework and can reunite the three Rio conventions. While upscaling will require context-specific adaptation, we believe community-led small-scale grazing exclusions to be a model system that can boost dryland restoration efforts. Overcoming current financial barriers – especially by engaging funding sources by both public and private sectors and improving monitoring frameworks – will be essential to unlock broader implementation. As global restoration efforts gain momentum, small-scale, community-led natural regeneration initiatives like grazing exclusions must be recognized and supported as critical building blocks for a more resilient and equitable future.

## Material and methods

### Study Area

This study was conducted in central Burkina Faso, within the administrative regions of Centre, Centre-Ouest, and Plateau Central (Figure S1). This area lies in the Sudano-Sahelian transition ecological zone, characterized by semi-arid woody savannah. The climate includes a short rainy season (July–October, 500–800 mm/year) and a prolonged dry season (November–June). This region exemplifies broader trends occurring across many Sahelian and sub-Saharan African countries, where rapid population growth (2.23% in Burkina Faso and 2.46% sub-Saharan Africa; World Population Prospects 2024 (www.population.un.org)) is intensifying pressure on natural resources. Projected increases in livestock farming and deforestation are likely to exacerbate existing pressures on ecosystems, contributing to accelerated land degradation. Most of Burkina Faso’s population lives in rural areas (67% in 2024; World Bank (www.data.worldbank.org)), relying primarily on small-scale subsistence farming. Around 80% of rural households keep domestic livestock (50), highlighting the critical role of land-use strategies like grazing exclusions in supporting both ecological restoration and local livelihoods.

### Study Sites

Since 2003, the international organization newTree (www.newtree.org) and its local partner, the association tiipaalga (www.tiipaalga.org), have established over 450 small-scale grazing exclusions (locally known in French as mise en défens) across Burkina Faso. Each grazing exclusion spans approximately 3 ha, totaling over 1,200 ha of restored land. These grazing exclusions are designed to exclude free-ranging domestic livestock, allowing native vegetation to regenerate naturally and degraded ecosystems to recover. As large wild mammals are absent from the study region no negative impacts on local fauna are expected. Grazing exclusions are implemented at the request of landowners, who retain full ownership and manage them with technical support from tiipaalga. Site selection follows a participatory process involving landowners, local communities, and tiipaalga, ensuring clear land tenure and alignment with local socio-cultural contexts (see Marcacci et al., 2025 (29) for more details). Only degraded lands can be selected as sites for a grazing exclusion.

Due to the current security situation, only grazing exclusions from central Burkina Faso were accessible. We selected 27 grazing exclusions as study sites, ensuring a minimum distance of 1 kilometer between any two of them to maintain their independence. As it can take a few years for the natural vegetation to recover, we excluded grazing exclusions younger than 6 years after consultation with experts from tiipaalga (see also Marcacci et al., 2025 (29)). The landscape around grazing exclusions is dominated by degraded wooded savannah interspersed with small-scale subsistence farms, villages, and widespread livestock grazing. Verbal consent was obtained from all landowners prior to fieldwork.

### Biodiversity Surveys

Vegetation and bird surveys, two good indicator groups to assess restored habitats, were conducted in all 27 grazing exclusions and 27 paired control sites. A pair of grazing exclusion and control site forms a “landscape”. Each landscape was relatively homogenous in terms environmental (including soil or topographic) and socioeconomic conditions. While grazing exclusion sites remained consistent across surveys, control sites differed between vegetation and bird surveys due to differences in spatial response scales (Figure S1).

Tree inventories were conducted within circular plots with a 50 m radius (∼ 0.78 ha), representing roughly one quarter of each grazing exclusion. One plot was placed inside each exclusion, at least 60 m from the fence to avoid edge effects. Each was paired with a control plot of the same size, located directly north of the exclusion with its center 100 m from the fence as shown in Figure S1. Within these plots, all trees with a diameter at breast height (DBH, measured at 130 cm) ≥ 5 cm were identified to species level, counted, and measured for DBH. Tree inventories were conducted between May and December 2023 by a local expert (Roland R. Kaboré). Additionally, within a 10 m radius subplot, herbaceous cover and mean height were estimated, including separate assessments for the dominant species. See Marcacci et al., (2025) (29) for more details.

Bird surveys were conducted using Passive Acoustic Monitoring (PAM) with Autonomous Recording Units (ARUs; AudioMoth Dev 1.2.0 (51)) during two seasonal periods: October–November 2022 (wet season) and March–April 2023 (dry season). One ARU was installed at the center of each grazing exclusion and a paired control site with similar vegetation cover, selected via remote sensing and field validation (Figure S1). Control sites were located on average 295 m (SD = 92 m) from grazing exclusions to ensure independence of acoustic recordings. Each ARU recorded for 2 hours per day (1 minute every 6 minutes), starting 30 minutes before sunrise, yielding 20 * 1-minute recordings per site per season (36 hours total across 54 sites). Due to limited coverage of African birds in AI models such as BirdNET, all recordings were manually annotated and expert-validated using Raven Lite (52) and ecoSound-web (53). Species richness was calculated as the total number of unique species detected per site across both seasons. See Quintas, Marcacci et al., (2025) (30) for more details.

### Socioeconomic Surveys

From May to December 2023, 54 individual face-to-face interviews were conducted with the heads of 27 farming households managing a grazing exclusion, as well as one neighboring household per site serving as a control (i.e., managing the area where control biodiversity data were collected), thus allowing to link socioeconomic to biodiversity data. All households were of comparable socioeconomic status and practiced mixed farming (crops and livestock). The interviews aimed to (1) document local uses of plant species for ecosystem service provisioning and (2) assess household income sources, with a focus on direct economic benefits derived from grazing exclusions. Respondents reported income from crop production, livestock production, non-agricultural activities, and resource-based products from grazing exclusions (e.g., shea butter, honey or fodder sales). Direct non-market benefits (e.g., own consumption of natural products) and indirect benefits from grazing exclusions (e.g., own use of fodder) could not be monetarily quantified, as households did not systematically track them – making our estimates of their contribution to annual income conservative.

To ensure ethical standards, all participants were informed of the study’s purpose and gave verbal consent prior to participation (written consent was not feasible due to illiteracy in some cases). Participation was voluntary and could be withdrawn at any time. Data were anonymized to protect participant identity. Interviews were conducted in the local language Mooré, based on a questionnaire originally written in French.

### Ecosystem Services and ES-Multifunctionality

All ecosystem services were quantified using ecological indicators derived from biodiversity and socioeconomic surveys following (35, 54). Because the goal was to evaluate the direct contribution of ecosystem services to farming households’ annual income, we focused exclusively on provisioning services. Socioeconomic interviews provided information on how each tree and herbaceous plant species was used locally. This information was then used to assign specific provisioning services to plant species (35). For each site, we quantified service provision by summing the number of trees associated with a given use (e.g., firewood, construction material, medicinal use, or non-timber forest products). For instance, the provision of non-timber forest products was estimated as the total number of trees identified as sources of such products. Fodder provision was estimated by calculating the biovolume of herbaceous species consumed by livestock, derived from field measurements of their percent cover and mean height. This metric is provided in m^3^·m^-2^. Information on indicators and calculations of each ecosystem services can be found in Table S2.

To capture the simultaneous delivery of multiple ecosystem services, we calculated ecosystem service multifunctionality (ES-multifunctionality), which integrates multiple service indicators into a single metric. This composite measure reflects the extent to which biodiversity supports the simultaneous provision of multiple ecosystem services (55). We used both the average and threshold approaches to quantify ES-multifunctionality. The average approach calculates the mean value across all standardized ecosystem service indicators for each site. In contrast, the threshold approach defines services as “delivered” when their value exceeds a predefined threshold, expressed as a percentage of the maximum observed value (56–57). To address the arbitrariness of selecting a single threshold, we used a multi-threshold approach, assessing service delivery across a gradient of thresholds from 1% to 99% (33). To reduce the influence of outliers in defining the maximum observed value, we calculated it as the average of the five highest values recorded across all sites (58). All statistical analyses were performed for both the average-based and threshold-based multifunctionality metrics.

### Carbon Stock and CO_2_ Sequestration

We estimated above- and below-ground carbon stocks and associated CO_2_ sequestration using biomass values derived from vegetation surveys (58). For trees, we applied species-specific or functional group-specific allometric equations developed for Burkina Faso to calculate wood volume and above and belowground biomass (Supporting text in Supporting Information), from which we derived Carbon stocks and CO_2_ sequestration estimates. Similarly, for herbaceous vegetation, we used allometric models tailored to semi-arid West African savannas, including Burkina Faso, to estimate both above- and below-ground biomass, and subsequently calculated corresponding carbon stocks and CO_2_ sequestration (59).

Total site-level Carbon storage and sequestration were calculated by summing contributions from both tree and herbaceous biomass. As our surveys excluded trees with a diameter at breast height (DBH) < 5 cm, our estimates are conservative, omitting Carbon stored in younger regenerating individuals. To complement these estimates and capture broader patterns of vegetation productivity, we also used the Normalized Difference Vegetation Index (NDVI) derived from remote sensing as a proxy for vegetation productivity and, indirectly, for carbon sequestration (16). We used Sentinel-2 satellite imagery with a 10-m spatial resolution and extracted NDVI from the same 50-m-radius plots in which vegetation surveys were conducted within grazing exclusions. For control sites, we randomly generated five 50-m-radius plots within a 500-m radius of the center of grazing exclusions, but located outside exclusion boundaries. In total, we analyzed 35 Sentinel-2 images (12-24 images available per site, depending on their location), covering the full period of the tree inventories (May-December 2023). All images were filtered using Cloud Score+ using the *cs* band with a threshold of 0.5. All remaining cloud-free pixels were then averaged to the site level and used for subsequent analyses (60).

### Statistical analyses

We first assessed the multiple ecological and economic benefits provided by grazing exclusions. To do so, we analyzed the direct effect of grazing exclusion on biodiversity (tree richness, number of trees, bird richness), ecosystem services (carbon sequestration, NDVI, ES-multifunctionality) and annual income. We also assessed each of the six provisioning ecosystem services (apiculture, construction wood, firewood, fodder, medicinal plants, non-timber forest products) individually in separate models. We built a univariate (generalized) linear mixed-effects model (*lme4* R-package, 61) for each of the response variables with treatment (factor with two levels: control and grazing exclusion) as sole predictor and landscape units as random intercept to account for the paired design. We selected the best error distribution for each model based on model diagnostics and AICc values.

We then built a structural equation model (SEM) to investigate the direct and indirect effects of grazing exclusion, tree richness, number of trees and ES-multifunctionality on annual income. We used a method so-called generalized multilevel path analysis, or piecewise SEM, which allows to test causal relationships with a relatively low sample size and to use a large variety of response distributions (R-package *piecewiseSEM,* 62). We first built a hypothetical SEM by combining three (generalized) linear mixed-effects models. Because tree richness and the number of trees were highly correlated, we built separate hypothetical SEM for each of these two variables. The first model tested the effect of grazing exclusion and time (since the establishment of the grazing exclusion) on tree richness/abundance to control for age effects. The second model tested the effect of grazing exclusion and tree richness/abundance on ES-multifunctionality. And the third model tested the effect of grazing exclusion and ES-multifunctionality on annual income. In all models, landscape unit was set as random intercept to account for the paired design. The best error distribution and model structure were selected prior to building the SEMs. We used Shipley’s d-separation test to detect missing paths and we assessed the goodness-of-fit of the SEMs with Fisher’s *C* statistics (63). We standardized models’ estimates using two SDs to improve the direct comparison of effect sizes between continuous and binary (here control *vs* grazing exclusion) predictors (64) using the *parameters* R-package (65).

Marginal and conditional R^2^ of all models were calculated with the *performance* R-package (66). The diagnostics of each individual model, including spatial autocorrelation of the residuals, were verified with the *DHARMa* R-package (67) and we used the variance inflation factor (VIF) to check for collinearity with the *CAR* R-package (68). All statistical analyses were conducted in R version 4.4.1 (69).

## Acknowledgements

We warmly thank all the staff of tiipaalga and newTree for their great work and logistical support as well as all farmers and landowners who allowed us to work on their land. We thank Ingo Grass, Clara Zemp and Sara Löfqvist for their comments on earlier versions of the manuscript and Liv Fritsche for her help with the remote sensing analyses.

## Supporting Information

### Supporting text

#### Carbon Stock and CO_2_ Sequestration estimation

##### Vegetation surveys

We estimated above- and below-ground carbon stock and associated CO_2_ sequestration using biomass values derived from vegetation surveys. Tree inventories were conducted within circular plots with a 50 m radius (∼ 0.78 ha). Within these plots, all trees with a diameter at breast height (DBH, measured at 130 cm) ≥ 5 cm were identified to species level, counted, and measured for DBH. Herbaceous cover and mean height were estimated within a 10 m radius subplot.

##### Tree biomass estimation

The volume of standing timber was obtained using allometric equations developed as part of the second national forest inventory (IFN2, 2018). For multi-stemmed trees, their equivalent diameters were calculated before applying the allometric equations to estimate the corresponding timber volume. The allometric equations are as follows:

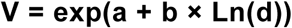

Where **V** is the volume of standing timber, **a** and **b** species or group-specific constants (see IFN2, 2018) and **d** tree diameter at breast height (130 cm).

Then, tree aboveground biomass (AGB) was estimated using:

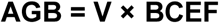

Where **V** is the volume of standing timber and **BCEF** is the biomass conversion and expansion factor, i.e. 2.11 for tropical dry forests (FAO, 2020).

As for belowground biomass (BGB), it is obtained by multiplying the value of above-ground biomass (AGB) by a coefficient **R**:

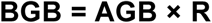

Where **R** = Stem-to-root ratio, which is 0.28 for tropical dry (FAO, 2020). Total tree biomass is the sum of above and belowground biomass:

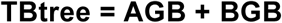

##### Herbaceous vegetation biomass

Herbaceous vegetation aboveground biomass was estimated using an allometric equation developed for West African semi-arid savannah (Savadogo et al., 2007):

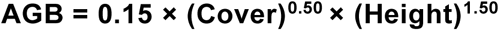

Where **Cover** is the fractional vegetation cover (0-1) and Height is the mean vegetation height (m). The equation yields in t ha^-1^. Biomass per plot was obtained by multiplying the per-hectare estimate by the plot area (0.0314 ha).

Belowground biomass was estimated using biome-specific root:shoot ratios following Mokany et al., (2006). For semi-arid savannahs, we applied a conservative ratio 1.5:1:

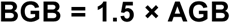

Total herbaceous vegetation biomass (TB) was calculated as:

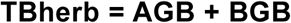

##### Carbon stock estimation

Total biomass (trees + herbaceous vegetation) was summed at the plot level and expressed per hectare Carbon stock (**C**) was calculated by applying the IPCC default carbon fraction of 0.47 (dry biomass carbon content) (IPCC, 2006)

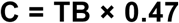

##### CO2 sequestration estimation

To convert carbon stock into CO_2_ equivalents (**CO_2_-e**), we multiplied Carbon stock by the molecular weight ratio of CO_2_ to carbon (44/12):

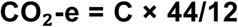

This yields the mass of atmospheric CO_2_ sequestered in vegetation biomass, expressed in t CO2 ha^-1^.

**Figure S1.**
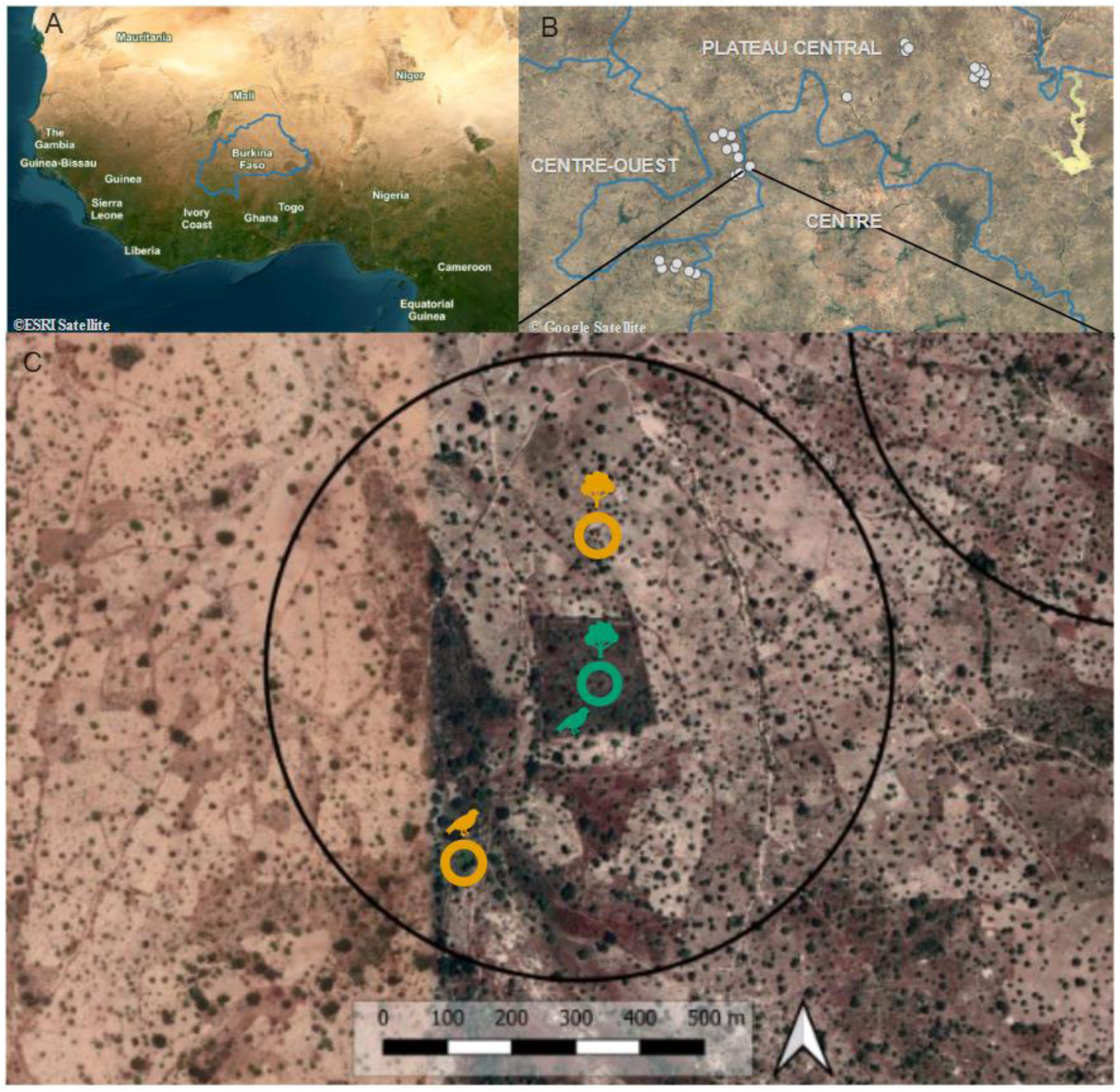
Location of Burkina Faso in West Africa (A). Locations of the 27 landscapes which are displayed with light grey dots (B). (C) Example of a landscape (black circle) with the grazing exclusion in the middle (green circle), and control sites (orange circles). Tree and bird icons correspond to where vegetation and bird surveys were conducted.

**Figure S2.**
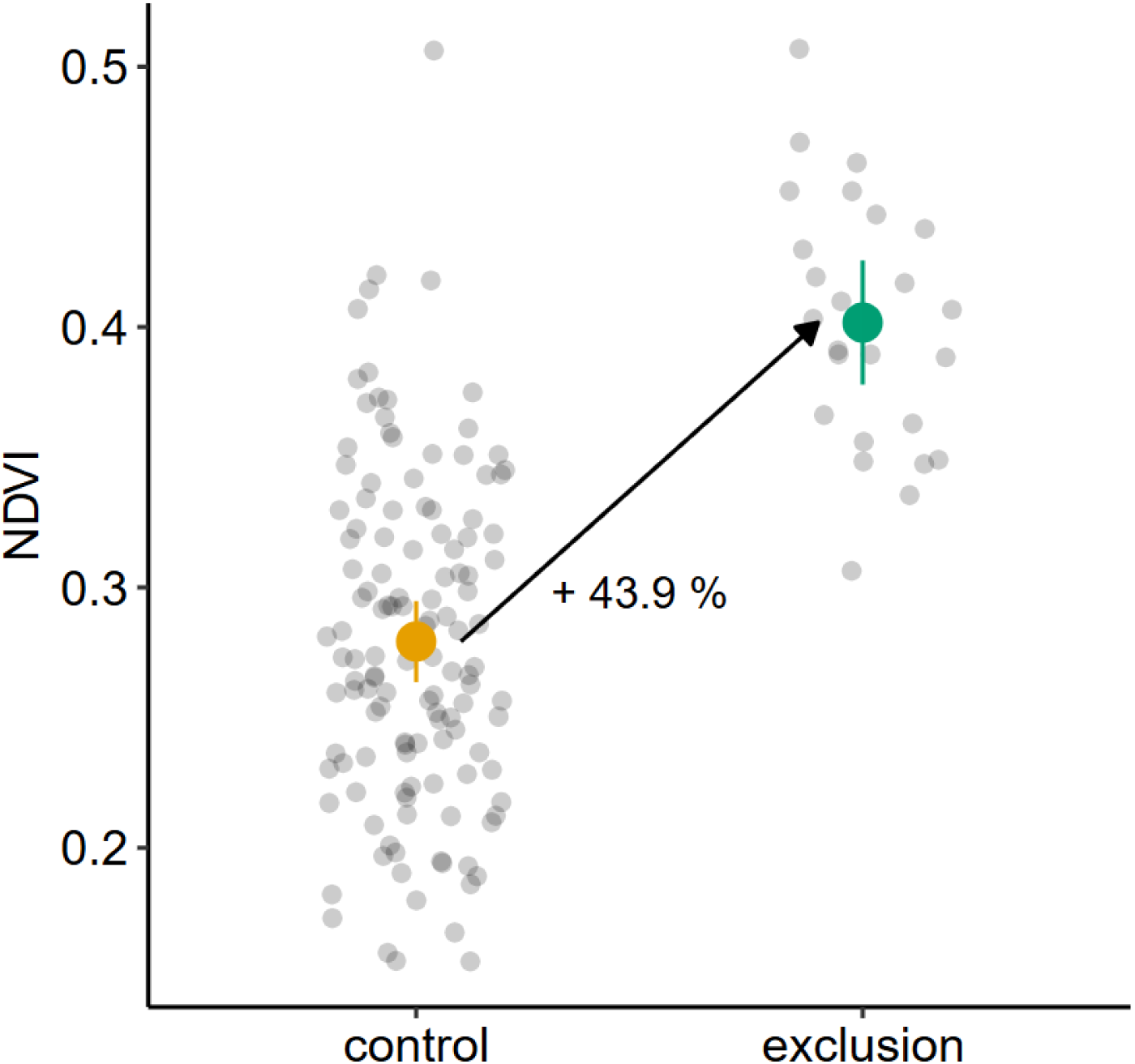
Positive effect of grazing exclusions on NDVI. Grey dots depict raw data, colored points (yellow for controls and green for grazing exclusions) the predicted means and confidence intervals drawn from a linear mixed-effects models.

**Figure S3.**
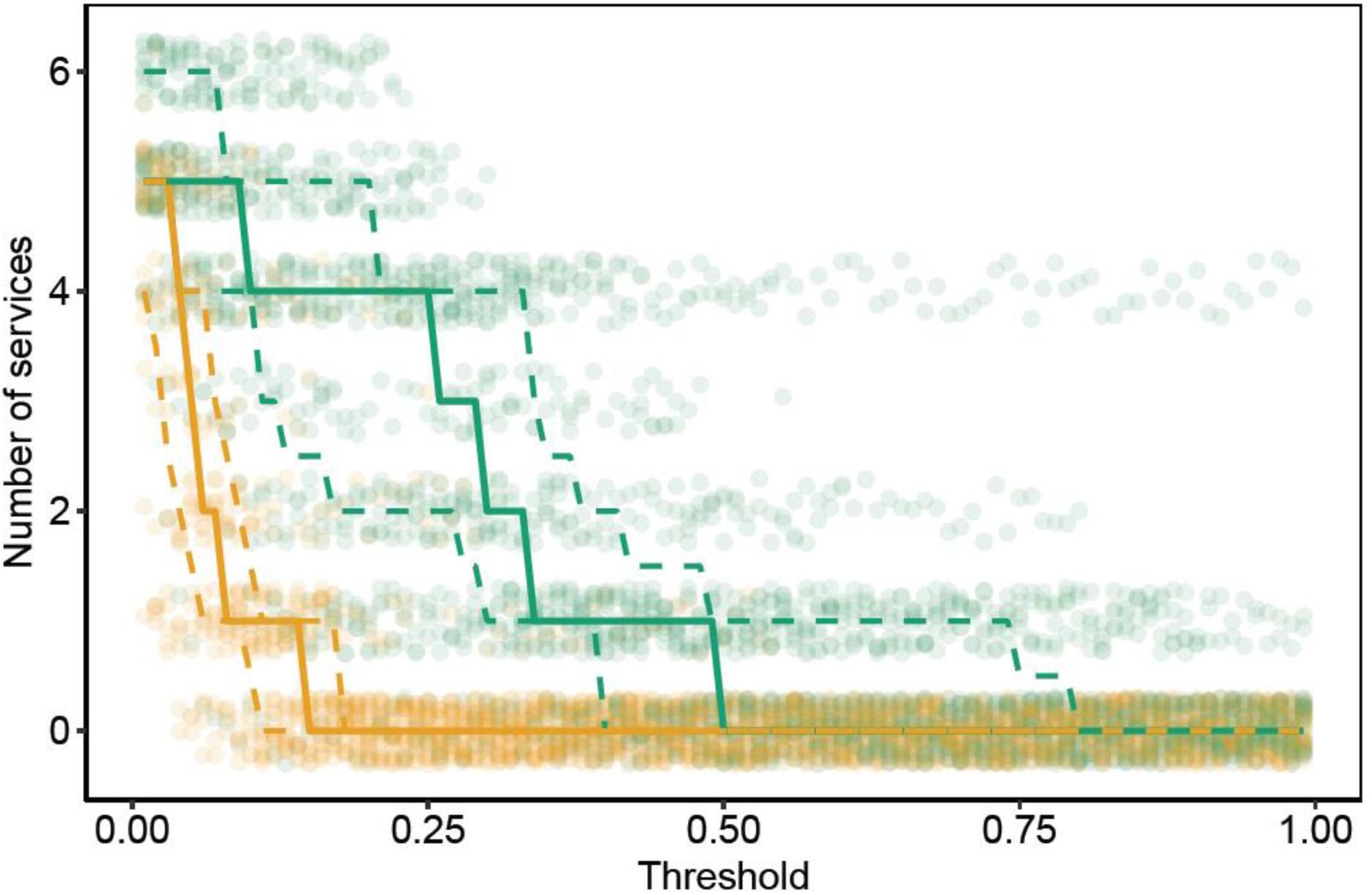
ES-Multifunctionality represents the number of ecosystem services that exceed a specific threshold, which is expressed as the percentage of the average of the five highest values recorded across all sites. Solid lines depict median values and dashed lines the quartiles (yellow for control and green for exclusion sites).

**Figure S4.**
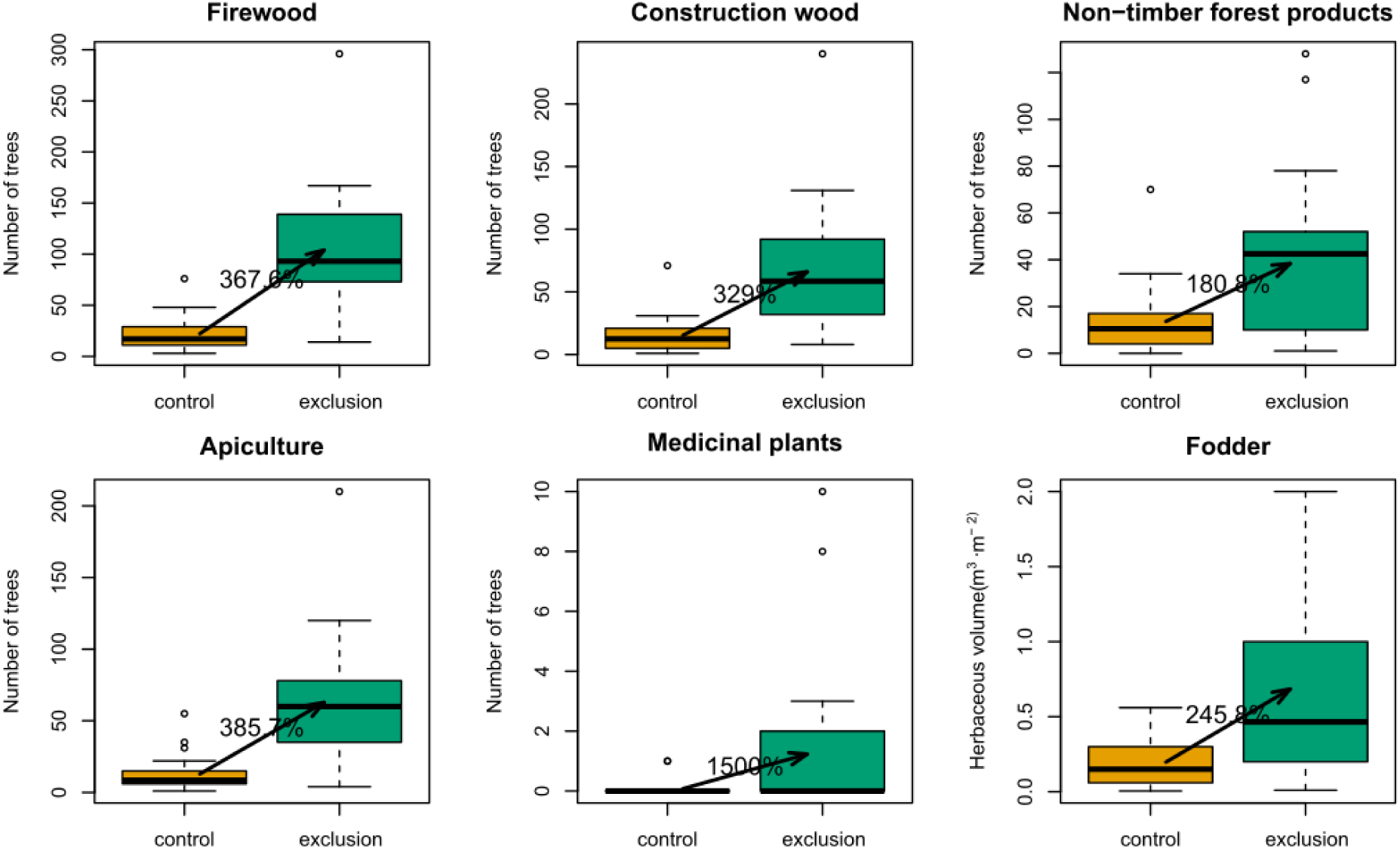
Positive effects of grazing exclusions on individual ecosystem services.

**Figure S5.**
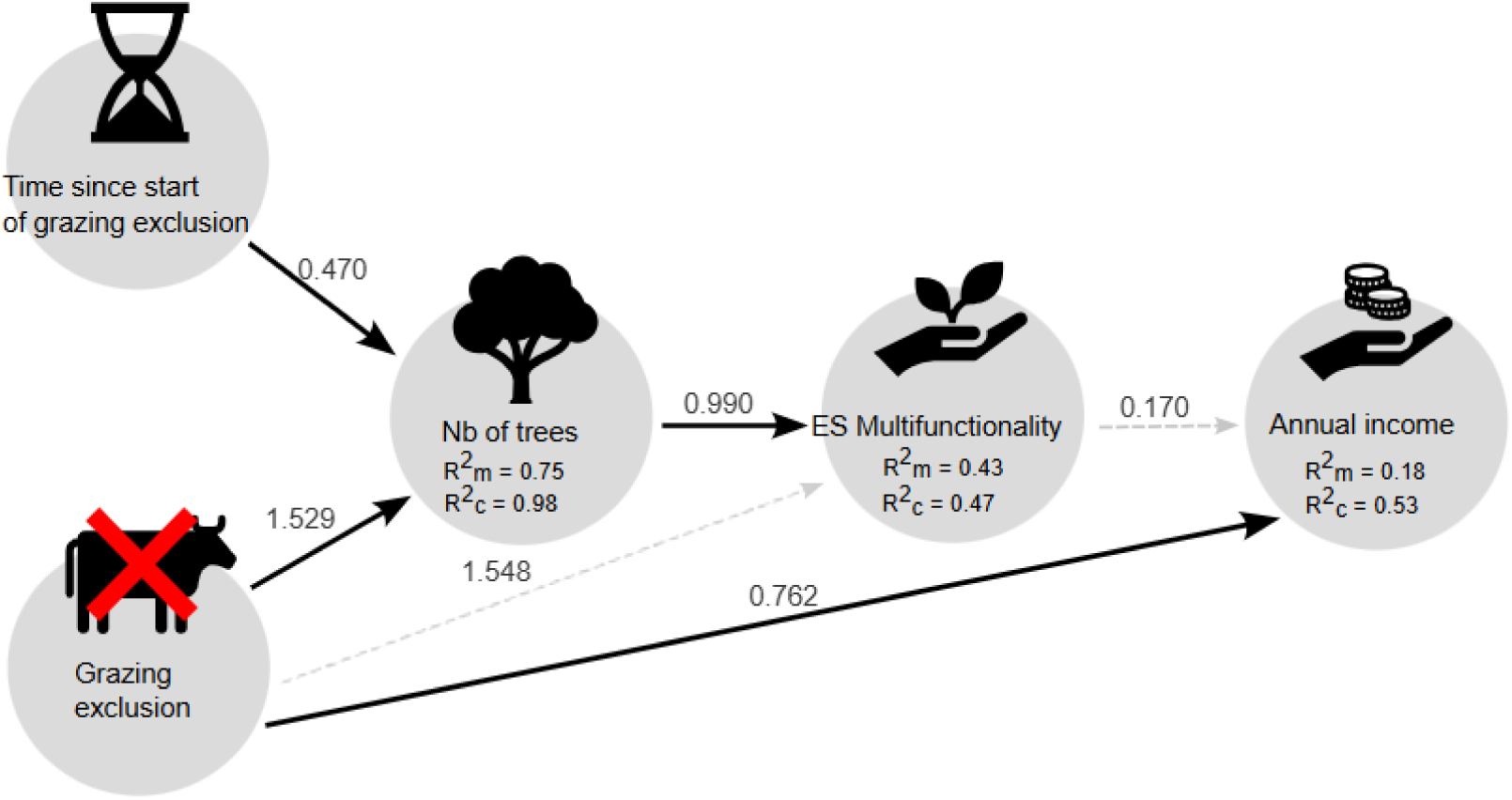
Path diagram illustrating the results of a Structural Equation Model investigating the direct and indirect effects of grazing exclusions, time, tree abundance and ES-multifunctionality on household income. Significant (p-value < 0.05) causal relationships are displayed with black solid arrows and non-significant relationships with grey dashed arrows. Standardized models’ coefficients are displayed next to the arrows and marginal and conditional R^2^ of individual model below the response variables.

**Table S1.** List of the provisioning ecosystem services measured with their ecological indicators and description.

| Ecosystem service | Indicator of ecosystem service provision | Description |
| --- | --- | --- |
| Apiculture / honey | Number of honey trees | Honey has a high market value and its production via sustainable apiculture can significantly contribute to household income. Due to landscape degradation, there is a lack of flowering resources for honeybees making the regeneration of honey trees critical for apiculture and honey production. |
| Construction wood | Number of trees used for construction | Based on tree species used to build constructions. |
| Firewood | Number of trees used as firewood | Almost all trees can be used as firewood, but some are not used due to their high value for other usage. In grazing exclusions, only wood from pruned trees is used for firewood. |
| Fodder | Volume of herbaceous plants consumed by livestock | Herbaceous plants such as *Andropogon gayanus* are used as fodder for livestock. They can be manually harvested (mown) inside grazing exclusions. |
| Medicine | Number of trees used in traditional medicine | Many trees are valued for their medicinal properties and are used for traditional medicine by local communities. |
| Non-timber forest products (NTFP) | Number of trees from which NTFP are derived | NTFP are natural products harvested from certain trees such as fruits and nuts. Some trees can directly be consumed while others such as shea (*Vitellaria paradoxa*) are processed into shea butter. Some of these products such as shea butter have a high market value and can contribute significantly to household income. |

**Table S2.**
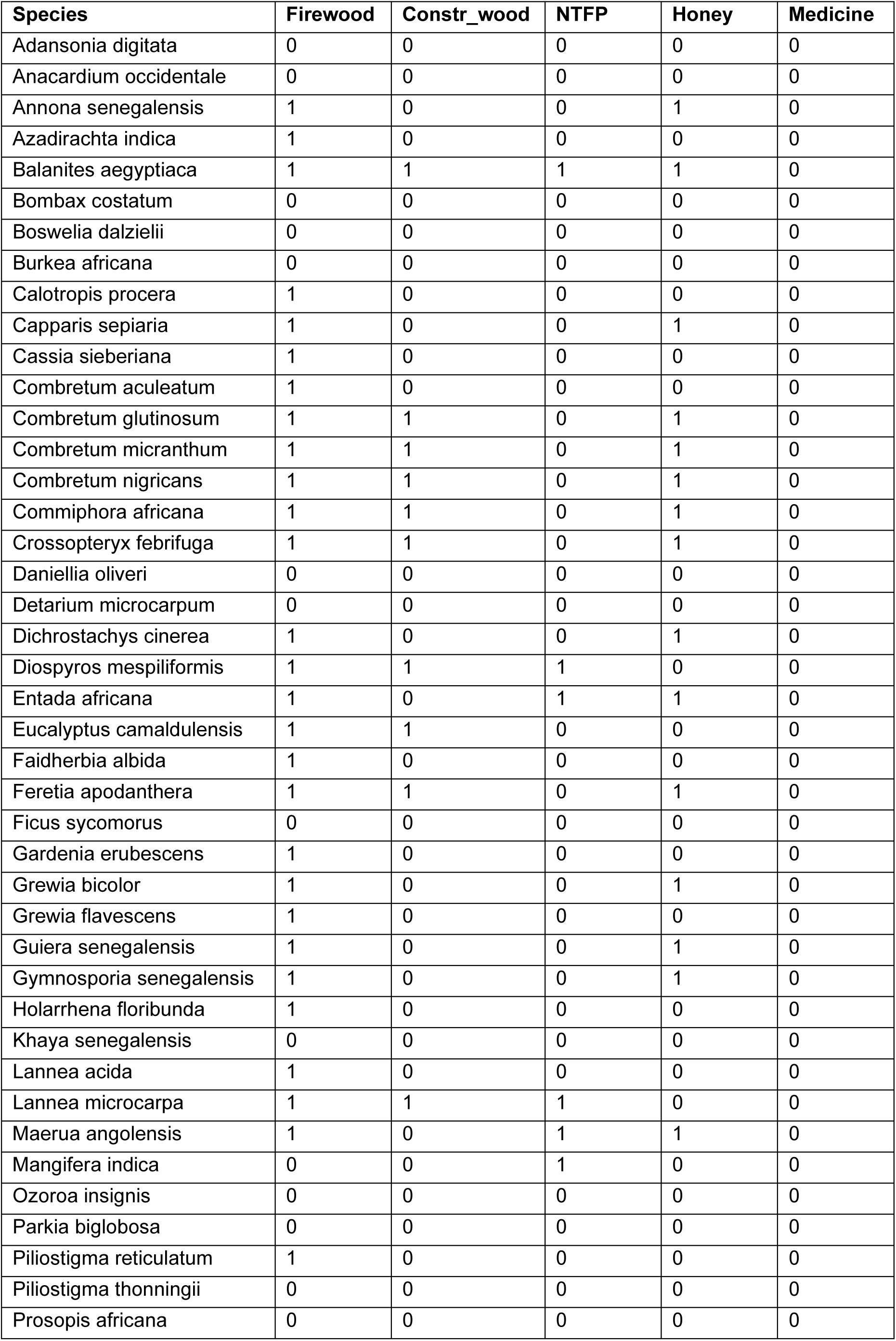
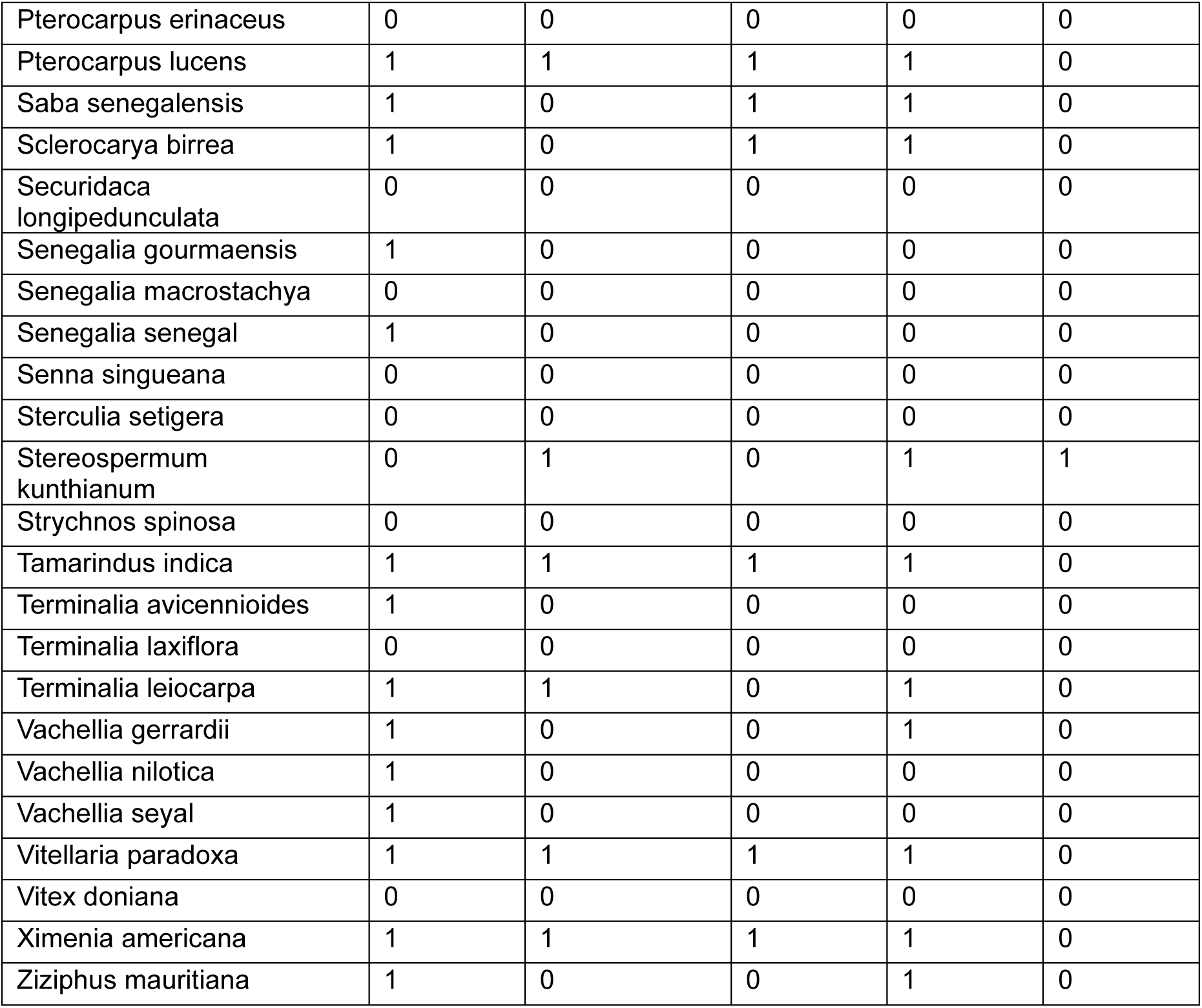
List of tree species recorded and their use (=1) by local communities.

**Table S3.** Coefficients of models testing the direct effect of grazing exclusions. Numbers indicate direction and magnitude of coefficient estimates, with the standard errors in brackets. Numbers in bold indicate p-value < 0.05.

| Response | Distribution | Intercept | Treatment | R<sup>2</sup><sub>c</sub> | R<sup>2</sup><sub>m</sub> |
| --- | --- | --- | --- | --- | --- |
| Tree richness | Poisson | **1.982**<br>(0.080) | **0.800**<br>(0.087) | 0.698 | 0.607 |
| Number of trees | Poisson | **3.096**<br>(0.099) | **1.529**<br>(0.044) | 0.982 | 0.726 |
| Bird richness | Gaussian | **29.039**<br>(0.774) | **5.810**<br>(0.757) | 0.692 | 0.356 |
| CO<sub>2</sub> sequestration | Gaussian | **131.16**<br>(16.37) | 24.03<br>(22.32) | 0.089 | 0.021 |
| Multifunctionality<br>(averaged) | Gaussian | **0.069**<br>(0.017) | **0.195**<br>(0.019) | 0.726 | 0.577 |
| Multifunctionality<br>(threshold = 50%) | Binomial | **-5.322**<br>(1.079) | **2.774**<br>(1.050) | 0.437 | 0.336 |
| Income | Gaussian | **982.38**<br>(223.83) | **1131.89**<br>(222.52) | 0.629 | 0.200 |
| NDVI | Gaussian | **0.28**<br>(0.008) | **0.12**<br>(0.011) | 0.538 | 0.356 |

**Table S.4.**
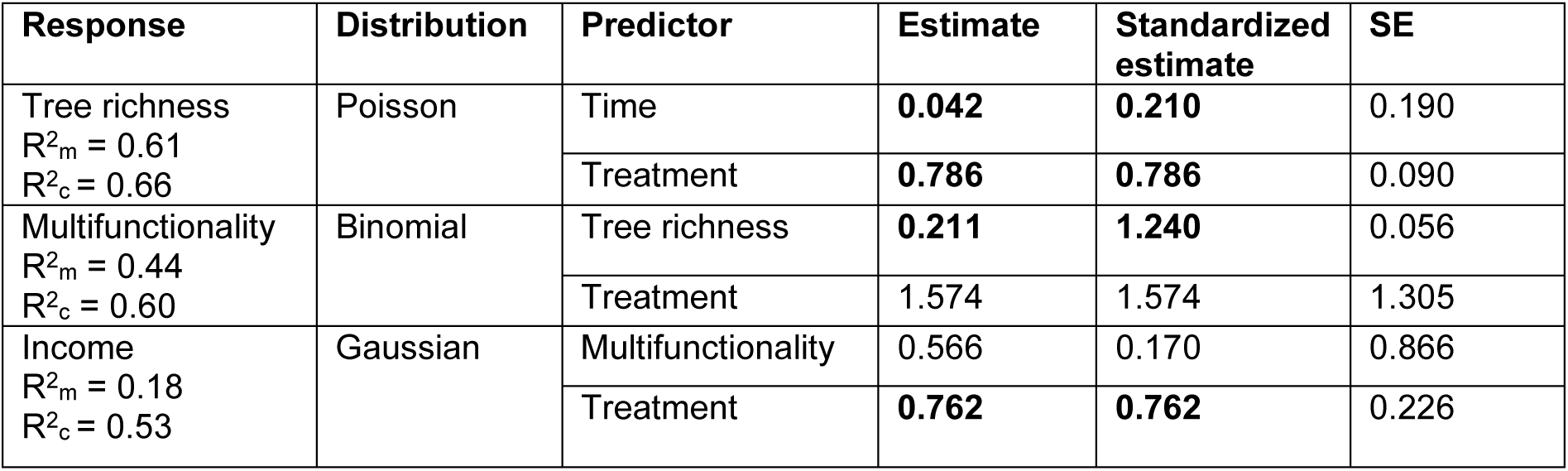
Coefficients of the SEM including tree richness (Fisher’s C = 5.722, p-value = 0.455). Standardized estimates were calculated with 2 SDs. Numbers in bold indicate p-value < 0.05.

| Response | Distribution | Predictor | Estimate | Standardized estimate | SE |
| --- | --- | --- | --- | --- | --- |
| Tree richness<br>$R^2_m = 0.61$<br>$R^2_c = 0.66$ | Poisson | Time | **0.042** | **0.210** | 0.190 |
|  |  | Treatment | **0.786** | **0.786** | 0.090 |
| Multifunctionality<br>$R^2_m = 0.44$<br>$R^2_c = 0.60$ | Binomial | Tree richness | **0.211** | **1.240** | 0.056 |
|  |  | Treatment | 1.574 | 1.574 | 1.305 |
| Income<br>$R^2_m = 0.18$<br>$R^2_c = 0.53$ | Gaussian | Multifunctionality | 0.566 | 0.170 | 0.866 |
|  |  | Treatment | **0.762** | **0.762** | 0.226 |

**Table S5.** Coefficients of the SEM including tree abundance (Fisher’s C = 6.342, p-value = 0.386). Standardized estimates were calculated with 2 SDs. Numbers in bold indicate p-value < 0.05.

| Response | Distribution | Predictor | Estimate | Standardized estimate | SE |
| --- | --- | --- | --- | --- | --- |
| Number of trees<br>$R^2_m = 0.75$<br>$R^2_c = 0.98$ | Poisson | Time | **0.095** | **0.470** | 0.037 |
|  |  | Treatment | **1.592** | **1.529** | 0.047 |
| Multifunctionality<br>$R^2_m = 0.43$<br>$R^2_c = 0.47$ | Binomial | Number of trees | **0.016** | **0.990** | 0.005 |
|  |  | Treatment | 1.548 | 1.548 | 1.305 |
| Income<br>$R^2_m = 0.18$<br>$R^2_c = 0.53$ | Gaussian | Multifunctionality | 0.566 | 0.170 | 0.866 |
|  |  | Treatment | **0.762** | **0.762** | 0.226 |

